# Reconciling *in vitro* and *in vivo* activities of engineered, LacI-based repressor proteins: Contributions of DNA looping and operator sequence variation

**DOI:** 10.1101/477893

**Authors:** Sudheer Tungtur, Kristen M. Schwingen, Joshua J. Riepe, Chamitha J. Weeramange, Liskin Swint-Kruse

## Abstract

One way to create new components for synthetic transcription circuits is to re-purpose naturally occurring transcription factor proteins and their cognate DNA operators. For the proteins, re-engineering can be accomplished via domain recombination (to create chimeric regulators) and/or amino acid substitutions. The resulting activities of new protein regulators are often assessed *in vitro* using a representative operator. However, when functioning *in vivo*, transcription factors can interact with multiple operators. We compared *in vivo* and *in vitro* results for two LacI-based transcription repressor proteins, their mutational variants, and four operator sequences. The two sets of repressor variants differed in their overall *in vivo* repression, even though their *in vitro* binding affinities for the primary operator spanned the same range. Here, we show that the offset can be explained by different abilities to simultaneously bind and “loop” two DNA operators. Further *in vitro* studies of the looping-competent repressors were carried out to measure binding to a secondary operator sequence. Surprisingly, binding to this operator was largely insensitive to amino acid changes in the repressor protein. *In vitro* experiments with additional operators and analyses of published data indicates that amino acid changes in these repressor proteins leads to complicated changes in ligand specificity. These results raise new considerations for engineering components of synthetic transcription circuits and – more broadly – illustrate difficulties encountered when trying to extrapolate information about specificity determinant positions among protein homologs.

## Introduction

As proteins are designed for biotechnological applications, one challenge can occur when *in vivo* outcomes do not match those of *in vitro* characterizations. We have encountered such an apparent discrepancy when making chimeras from the LacI/GalR transcription factors to be used in “logic gates” for bacterial computing [1-4].

Our design goal was to create repressor proteins that bound the same *lacO^1^* operator DNA sequence but responded to different small-molecule, allosteric ligands. To that end, chimeric repressors were created by joining the DNA binding domain of the *Escherichia coli* (*E. coli*) lactose repressor protein (“LacI”, UnitProtKP P03023) to the regulatory domains of paralogs, such as the *E. coli* purine repressor and galactose repressor proteins (respectively “PurR”, UnitProtKP P0ACP7; and “GalR, UnitProtKP P03024; Fig 1) [2, 5, 6]. The paralogous regulatory domains also mediated the dimerization needed to create a high affinity binding site for one DNA operator [7-9]. *In vivo* assays with these chimeras showed the desired outcomes: All chimeras repressed the natural *lac* operon and responded to the small molecule recognized by the paralogous regulatory domain (see Fig 2 in Meinhardt *et al.* [2]).

**Figure 1.**
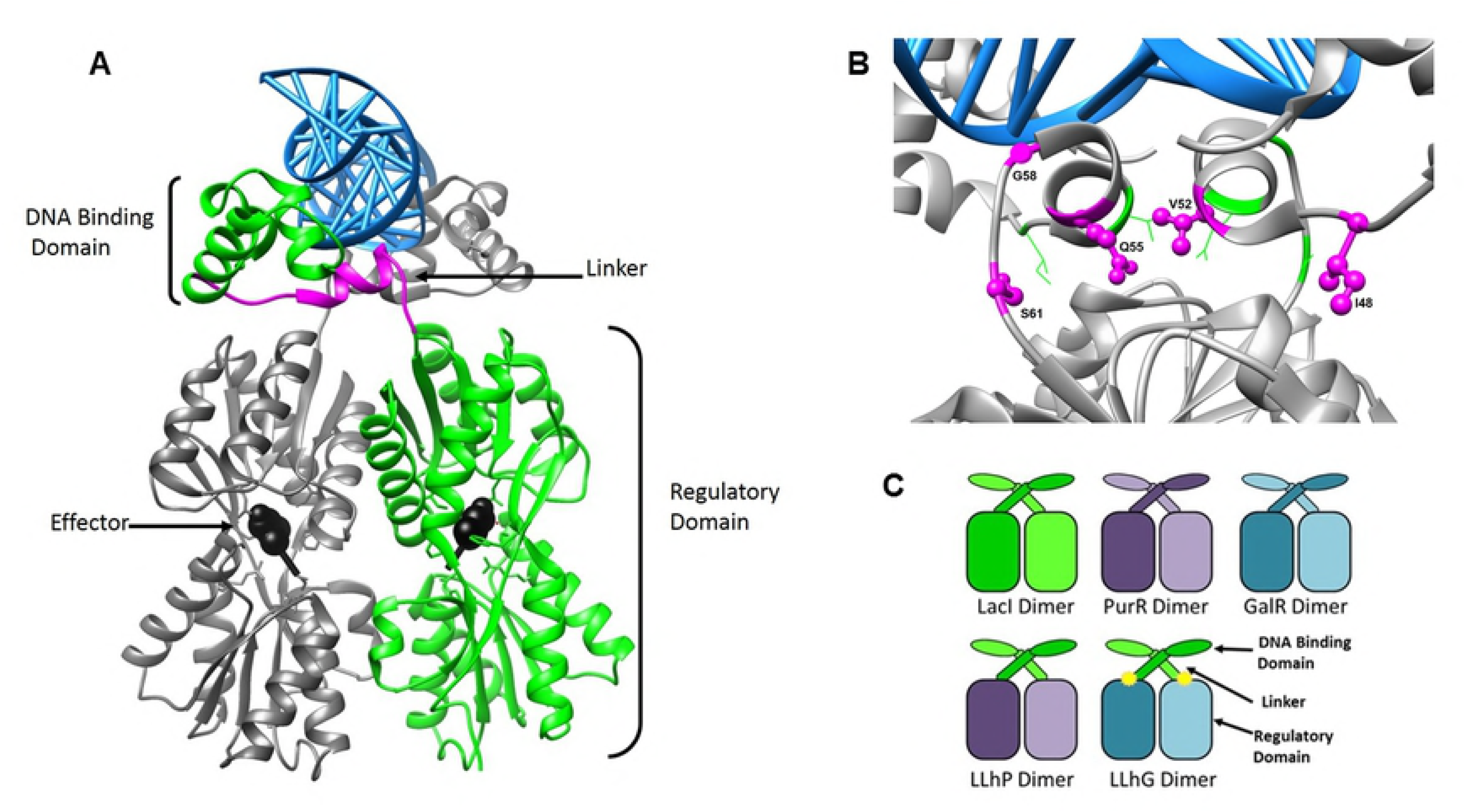
Ribbon and cartoon structures of LacI/GalR homologs. **(A)** The homodimer of the lactose repressor protein (LacI) (PDB ID 1EFA [7]) is shown with one subunit as a gray ribbon and the other in green. On the “green” monomer, the linker region is shown in magenta. The protein dimer is bound to DNA, which is depicted as a blue ladder. Allosteric effector is bound in the regulatory domain and represented as black spheres. The figure was rendered using UCSF Chimera [10]. **(B)** The LacI protein structure has been rotated and zoomed to show positions 48, 52, 55, 58 and 61 in the linker region. Amino acids nearest the plane of the viewer are shown in magenta ball-and-stick; those facing towards the rear of the structure (on the partner linker region) are in green wireframe. **(C)** The domain structure of the wild-type LacI homodimer is represented as a green cartoon; the PurR homodimer is represented in purple; and the GalR homodimer is represented in teal. These color schemes are used to indicate the source of the DNA binding domains (small ovals; LacI positions 1-44), linkers (bars; LacI positions 45-61), and regulatory domains (large ovals; PurR positions 60-340 or GalR positions 60-343) in the chimeric repressors “LLhP” and “LLhG”. All variants of LLhG in this manuscript contain the E62K mutation (“+K”), indicated by the yellow asterisk, as well as the “E230K” mutation (described in Materials and Methods).

**Figure 2.**
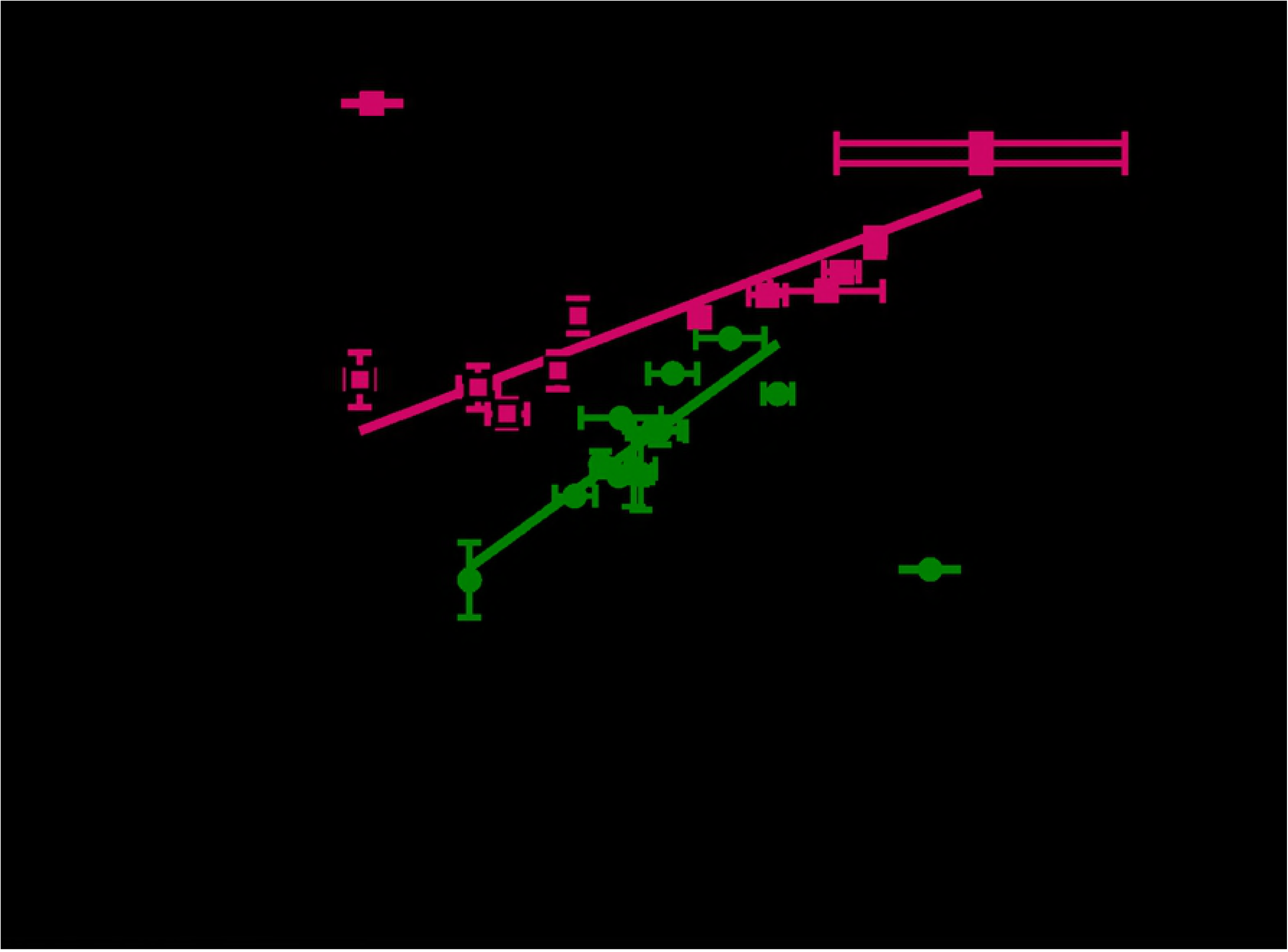
Comparison of LLhP and LLhG+K variants binding operator *lacO^1^*. *In vitro* binding to *lacO^1^* versus values from *in vivo* repression assays for variants of LLhP (magenta squares) and LLhG+K (green circles). Lines represent the best fit to the data and correlation coefficients are consistent with the linear relationship expected for these *in vivo* concentrations [3, 13]. Both X and Y error bars represent the standard deviations of averages determined from at least three separate experiments. Repression data were taken from [11]; for comparison among multiple chimeras, these published values were reported with a different normalization scale than the separate normalizations previously used for LLhP and LLhG+K in [5, 6]; error propagation was also revised. The arrows **outside the axes indicate that repression was enhanced as DNA binding affinity became tighter. In addition to altered** affinity from amino acid changes, LLhP had enhanced binding in the presence of 0.4 mM co-repressor hypoxanthine [4, 6]; values were determined +/- this effector; “plus” data are indicated with black-outlined squares.

Next, in exploring the outcomes that arose from amino acid changes in the interface between the DNA-binding and regulatory domains [5, 6, 11, 12], we purified sets of variants for the LacI:PurR chimera (“LLhP” [6]) and the LacI:GalR chimera (“LLhG+K” [5]) for biophysical studies [3, 4]. This allowed us to compare *in vivo* repression with *in vitro* DNA binding affinities for the primary operator of the *lac* operon (*lacO^1^*). Reassuringly, both sets of variants showed the expected relationship, which for the high protein concentrations in the *in vivo* assays should be linear [3, 13]. However, when the two studies were compared to each other, LLhP variants had weaker repression than LLhG+K variants, even though their *K*_d_ values for binding to the natural *lacO^1^* operator spanned the same range (Fig 2). This was unexpected: To our knowledge, the published literature about the *lac* operon, PurR, and GalR does *not* indicate that direct interactions with heteroproteins are expected to affect LLhG or LLhP repression in this setting. Thus, we explored the contributions that might arise from interactions of engineered repressors with alternative operators *in vivo* (Table 1).

**Table 1.**
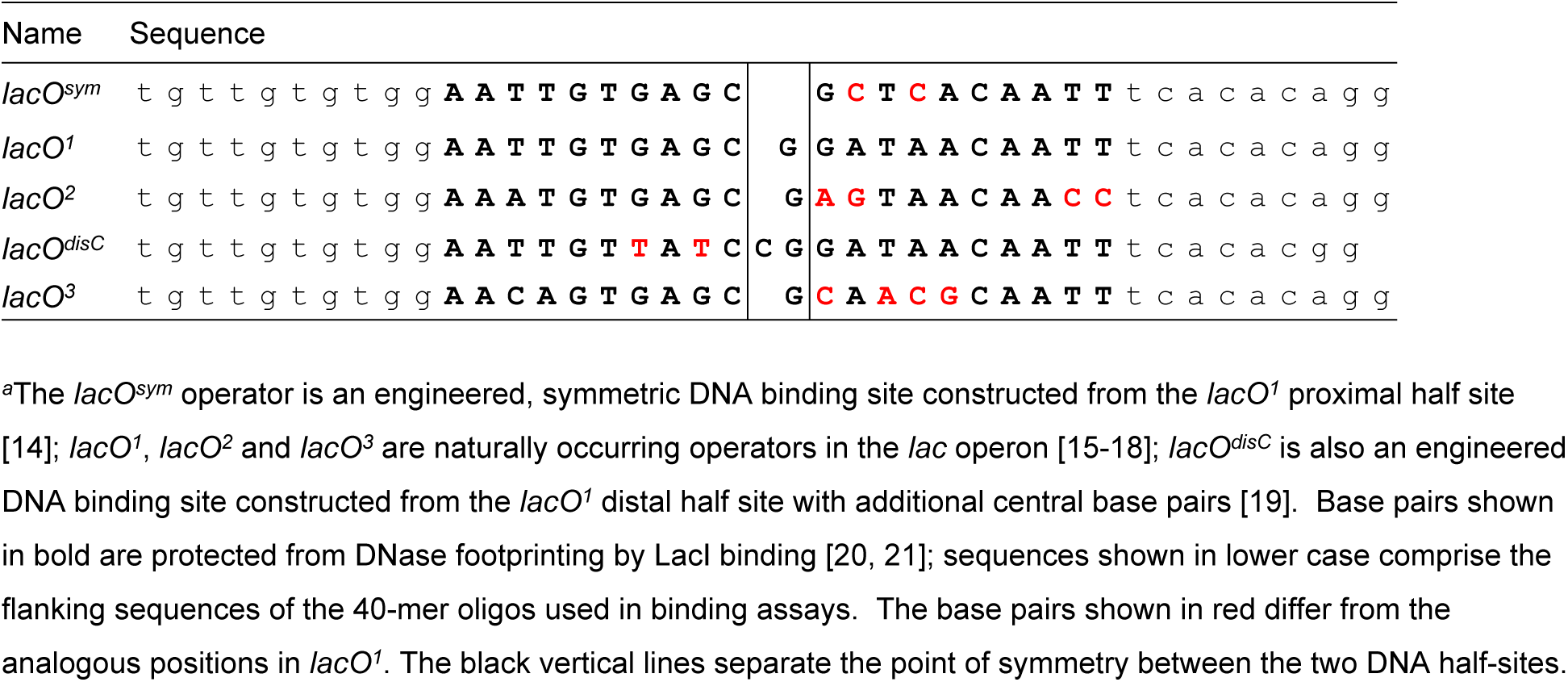
Relevant *lac o*perator sequencesa^a^

Here, we report that the tighter LLhG+K repression is consistent with this repressor protein looping two DNA operator sites, most likely *lacO^1^* and *lacO^2^*, that were present in the *in vivo* assays. *In vitro* experiments were also carried out to determine whether amino acid changes in the LLhG+K variants altered *K_d_* for *lacO^2^* in addition to the previously-measured changes in *lacO^1^ K_d_* values [3]. Surprisingly, binding to *lacO^2^* showed very little sensitivity to any of the amino acid variants tested in LLhG+K. To further assess the ligand specificity of these variants, additional experiments showed that binding to the tight-binding *lacO^sym^* operator was sensitive to the LLhG+K amino acid changes, whereas binding to the *lacO^disC^* operator was weaker than the limit of the assay. These unexpected changes in specificity raise new considerations for engineering components of synthetic transcription circuits and – more broadly – for extrapolating information about specificity determinant positions among protein homologs.

## Materials and Methods

### Ruling out “trivial” sources of repression differences

The discrepancy between LLhP and LLhG+K repression shown in Fig 2 could arise if LLhP variants were expressed at lower levels than the LLhG+K variants. However, *in vivo* protein concentrations were previously estimated to be >2500 copies per *E. coli* cell for all LLhP and LLhG+K variants [2, 11]. This is in vast excess over the single *lac* operon per genome, which makes it unlikely that differences in LLhP and LLhG+K repression are due to altered protein expression.

Another possible source of the discrepancy could be the *in vivo* presence of endogenous allosteric effectors. However, PurR is only known to have natural co-repressors – hypoxanthine and guanine – which enhance DNA binding/repression [22, 23]. Likewise, when surveyed with a variety of small molecules, LLhP repression only responded to the known PurR co-repressors and no gratuitous inducers have been identified to date [2, 4, 6]. In contrast, wild-type GalR responds to the natural inducer galactose and the gratuitous inducer fucose, which weaken DNA binding and repression [24]. Again, LLhG+K showed a similar response profile [2, 3, 5], and no gratuitous co-repressors have been identified to date. Thus, even if allosteric effectors were endogenous in the *in vivo* repression assays, their known influences are opposite to the discrepancy illustrated in Fig 2.

We also considered differences in the *in vitro* binding conditions of LLhG+K and LLhP. Binding affinities for LacI and LLhP variants were assayed in “FBB” buffer (10 mM Tris-HCl, pH 7.4, 150 mM KCl, 5% DMSO, 0.1 mM EDTA, and 0.3 mM DTT), but LLhG+K variants appeared to aggregate in this buffer over the course of the assay [3]. Relative to FBB, the successful LLhG+K binding buffer had a slightly lower pH, more reducing equivalents, and lacked DMSO (see below). However, LLhP DNA binding in the LLhG+K binding buffer produced essentially identical values to those previously reported [4]. Thus, the *in vitro* buffer differences were unlikely to be the source of the discrepancy illustrated in Fig 2.

### Proteins and purification

Plasmids expressing the coding regions of full-length LacI (plasmid numbers #31490 and #90058), LLhP (#90038), and LLhG (#90051) are available from addgene (https://www.addgene.org/). Variants of the LLhG/E62K protein (“LLhG+K”) were purified and DNA binding was carried out as described in Tungtur *et al.* [3]. This variant was previously chosen for mutagenesis because it repressed transcription more tightly that the parent “LLhG” chimera [5]. As before, all LLhG+K variants also carried the “E230K” mutation, which was required to alleviate bacterial toxicity [5]. Notably, DNA looping occurred in the parent LLhG+K chimera, despite the presence of the E230K substitution [2], which diminished looping in wild-type GalR [25]. Additional amino acid changes assessed in this study were located in the linker region of LLhG+K, as indicated in the figures and tables.

A brief description of LLhG+K purification is as follows: Variants were constitutively expressed from the plasmid pHG165a [5] and grown overnight in BLIM cells [26] in 2xYT media. Cell pellets were resuspended in cold breaking buffer (12mM Hepes, 200mM KCl, 1mM EDTA, 5% glycerol, 0.3 mM DTT, pH to 8.0) with 1 protease inhibitor tablet (ROCHE Diagnostics, Indianapolis, IN, USA) and frozen at −20°C. Following (i) cell lysis *via* freeze/thaw with lysozyme (Fisher Scientific) and DNA degradation via DNAse (Sigma-Aldrich Chemical Company), (ii) centrifugation, (iii) 37% ammonium sulfate precipitation and (iv) dialysis, the final purification step comprised a phosphocellulose (Whatman P-11) ion exchange column. LLhG+K proteins were eluted from the column using a linear gradient of Buffer A (12mM Hepes, 50mM KCl, 1mM EDTA, 5% glycerol, 0.3 mM DTT, pH to 8.0) and Buffer B (12mM Hepes, 500mM KCl, 1mM EDTA, 5% glycerol, 0.3 mM DTT, pH to 8.0). Protein elution occurred near conditions of 50% buffer A/50% buffer B. Aliquots of purified protein were stored at −80°C.

## DNA binding assays

Prior to DNA binding assays, purified LLhG+K variants required exchange into reducing conditions [3]. Protein variants were dialyzed against in HEPES/DTT buffer (12 mM Hepes, pH 7.53, 150 mM KCl, 0.1 mM EDTA buffer and 3 mM DTT) for 30 minutes in each of two buffer volumes; a third buffer exchange was into Tris/DTT buffer (10 mM Tris, pH 7.13, 150 mM KCl, 0.1 mM EDTA, and 3 mM DTT). The high concentrations of DTT precluded using A_280_ to determine concentrations of the LLhG+K variants. Therefore, protein concentration was estimated using the Bradford assay (BioRad, Inc., Hercules, CA), with bovine serum albumin (Fisher Biotech, Fair lawn, NJ, 07410) as a standard. In order to more precisely determine the concentration of protein competent for binding DNA, the activity of each protein preparation was determined by stoichiometric assays [27] to be between 70 and 99%. Activities were used to correct *K*_d_ values determined from binding titrations.

DNA binding affinities for LLhG+K and variants were measured by binding protein to 32P-labelled *lacO^2^*, *lacO^sym^*, and *lacO^disC^*. For most variants, *K_d_* values for *lacO^1^* were reported in [3]; binding data for a variant new to this work is shown in S5 Fig. All operator sequences (Table 1) comprised the central region of a 40 basepair, double-stranded DNA oligomer [28] and were synthesized by Integrated DNA Technology (Coralville, IA) and radiolabeled as in Zhan *et al.* [28]. After mixing protein and DNA, a 30-minute equilibration was allowed prior to filtration through nitrocellulose filter paper using a 96 well dot blot apparatus. Pseudo-equilibrium measurements were made by quickly separating the free and protein-bound DNA through nitrocellulose filter paper, which has been well-established for wild type LacI (*e.g.* [28]) and LLhP [4]. For affinity assays, the DNA concentration was fixed at least 10-fold below the value of *K*_d_ [27].

DNA binding affinities were determined in both the absence and presence of 10 mM inducer sugar fucose. Results were analyzed with nonlinear regression using the program GraphPad Prism 5 (GraphPad Software, Inc., La Jolla, CA) to determine values of *K*_d_, using:

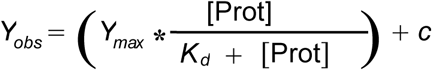

where “Y_obs_” is the observed signal from ^32^P-DNA, “Y_max_” is the signal observed at saturation, “[Prot]” is the concentration of the LLhG+K variants, “c” is baseline value of the ^32^P-DNA signal, and “*K*_d_” is the equilibrium dissociation constant.

Reported values in Table 2 are the average and standard deviation for at least three separate determinations, using at least two different protein preparations. Note that the values of standard deviations were larger than the errors of the fit.

**Table 2.**
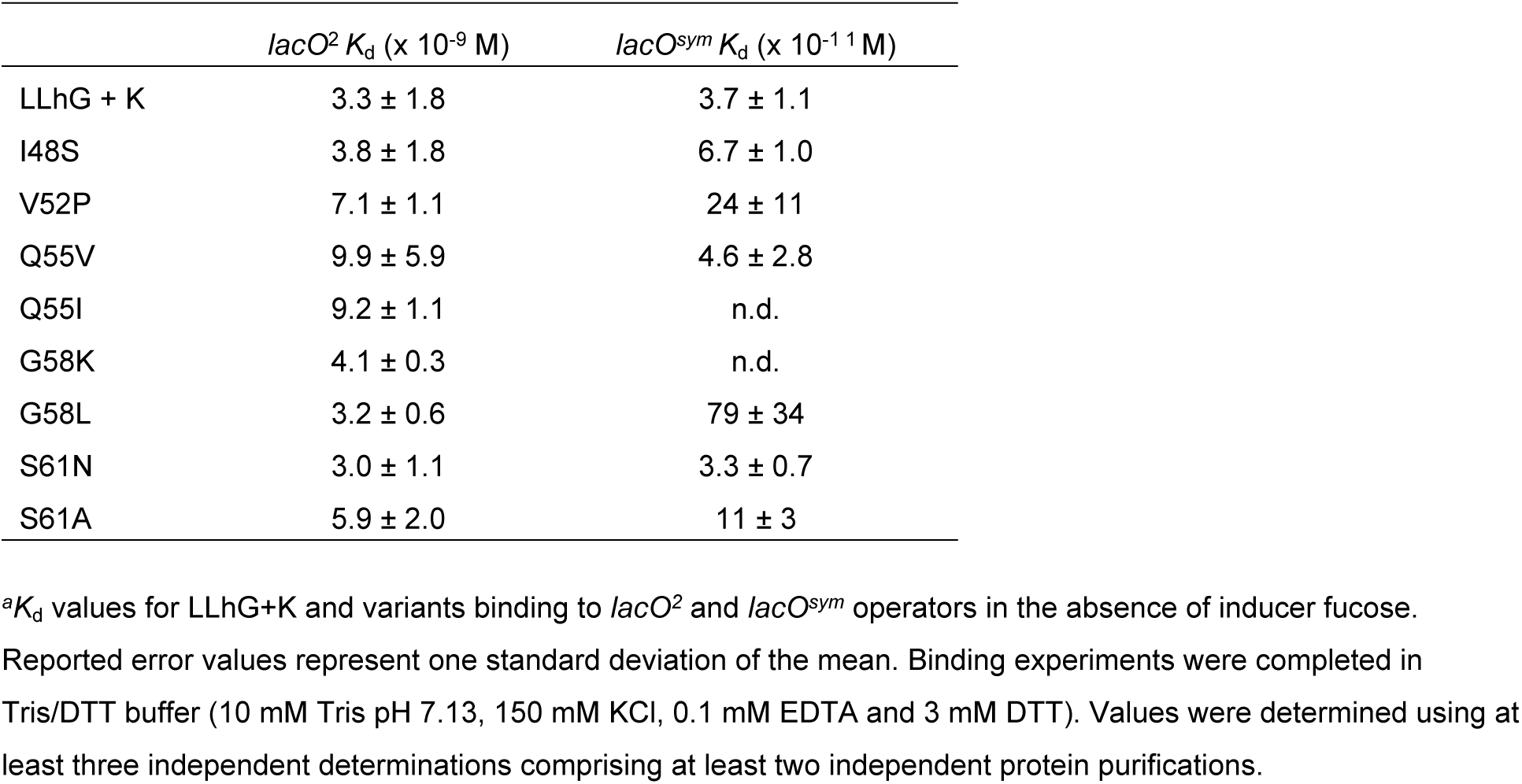
Operator binding by LLhG+K variants^a^

| | *lacO*<sup>2</sup> $K_d$ (x 10<sup>-9</sup> M) | *lacO*<sup>sym</sup> $K_d$ (x 10<sup>-11</sup> M) |
| --- | --- | --- |
| LLhG + K | 3.3 ± 1.8 | 3.7 ± 1.1 |
| I48S | 3.8 ± 1.8 | 6.7 ± 1.0 |
| V52P | 7.1 ± 1.1 | 24 ± 11 |
| Q55V | 9.9 ± 5.9 | 4.6 ± 2.8 |
| Q55I | 9.2 ± 1.1 | n.d. |
| G58K | 4.1 ± 0.3 | n.d. |
| G58L | 3.2 ± 0.6 | 79 ± 34 |
| S61N | 3.0 ± 1.1 | 3.3 ± 0.7 |
| S61A | 5.9 ± 2.0 | 11 ± 3 |

<sup>a</sup> $K_d$ values for LLhG+K and variants binding to *lacO*<sup>2</sup> and *lacO*<sup>sym</sup> operators in the absence of inducer fucose.
Reported error values represent one standard deviation of the mean. Binding experiments were completed in Tris/DTT buffer (10 mM Tris pH 7.13, 150 mM KCl, 0.1 mM EDTA and 3 mM DTT). Values were determined using at least three independent determinations comprising at least two independent protein purifications.

## Results

The stronger repression for LLhG+K variants, as compared to LLhP variants (Fig 2), could be explained by several phenomena. Thus, we first ruled out the “trivial” explanations of different protein expression levels and *in vitro* buffer conditions (see Materials and Methods). Next, we considered two other possible differences: Either the LLhP variants were competed away from the *lac* operator, or the local concentration of LLhG+K was enhanced by some mechanism. The first possibility could arise if LLhP variants showed tighter non-specific DNA binding than did LLhG+K variants. Although this remains a formal possibility, it would be very difficult to test since every base pair in a DNA sequence is the start of a distinct binding site, and different nonspecific DNA binding sequences can have different binding affinities [29-32].

In addition, we had prior experimental evidence [2] for the second option (enhanced LLhG+K local concentration). Local concentration can be increased when one protein (or protein complex) simultaneously binds two distal binding sites on DNA, “looping out” the intervening DNA sequence (Fig 3 A-C) [33]. Looping by LacI/GalR repressors requires tetramerization (since a homodimer is the unit for binding one DNA operator), and several homologs exhibit various tetramerization mechanisms.

**Figure 3.**
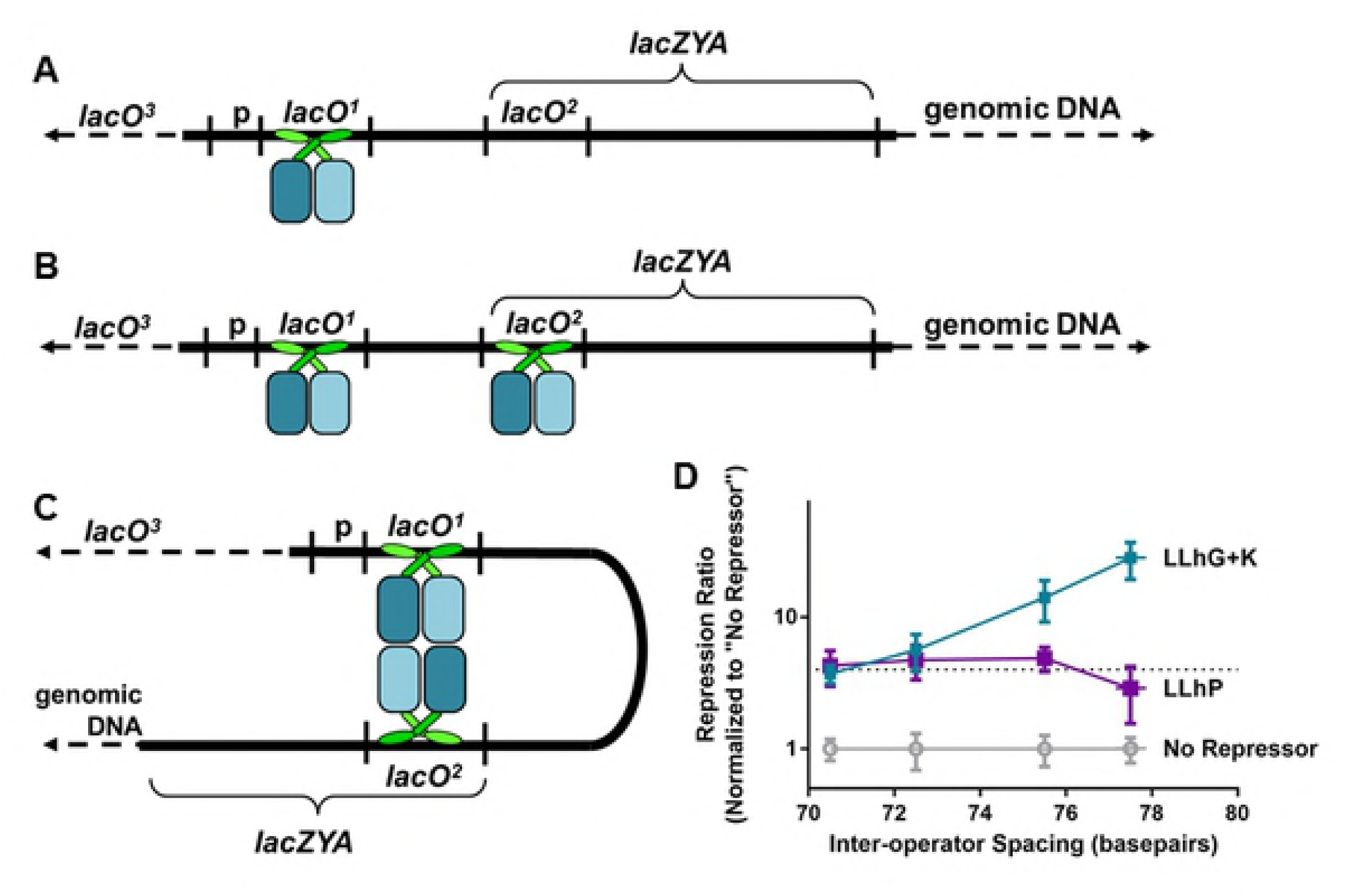
Looping in the *lac operon*. **(A)** When dimeric repressor is bound to the *lacO*^1^ DNA operator, transcription of the downstream *lacZYA* genes are repressed. **(B)** Dimeric repressor protein is capable of binding other sites in the *E. coli* genome such as *lacO*^2^, *lacO*^3^ and non-specific genomic DNA. **(C)** Tetrameric LacI can simultaneously bind two operator sites, leading to DNA looping. The regulatory domains of two wild-type GalR dimers also have the capability to form protein-protein interactions *via* its regulatory domains, which provides another means to facilitate tetramerization and DNA looping. **(D)** Prior experiments indicated that LLhG+K has looping capabilities, similar to its parent protein GalR [2]. Since DNA looping depends highly on inter-operator spacing (x-axis), *in vivo* repression can be altered by changing this distance. In the experiments shown, repression of the reporter gene was assessed using four strains of *E. coli*, containing a *lacZ* gene under control of the *lacO*^sym^ and *lacO*^2^ operators. Values were normalized to a “no repressor” control, and higher values represent increased repression. Note that LLhG+K repression was sensitive to operator spacing, whereas LLhP was not.

For example, LacI has an additional C-terminal tetramerization domain that mediates formation of a dimer-of-dimers [34-37] and can simultaneously bind two operators [33]. For wild-type LacI, looping enhanced *in vivo* repression ~50-fold [20, 33, 38-44]. In *in vitro* studies, LacI binding to DNA containing two operators had tighter affinity than expected from the sum of binding two, single operators [15]. In another example, full length GalR exhibited tetramerization when it participated in a “repressosome” complex with the hetero-protein “HU”. HU facilitated repressosome formation and looping *via* DNA bending; the repressosome complex facilitated and was stabilized *via* homomeric contacts between the regulatory domains of two GalR dimers [25, 45-50]. (Although GalR may directly interact with HU under some conditions [51], the heteroprotein interaction did not appear to occur in the repressosome and individual GalR dimers can repress transcription [52].) Notably, PurR and LLhP lack tetramerization domains; furthermore, no tetramerization has been observed to occur among the PurR or LLhP regulatory domains, even at high concentrations used in small angle X-ray scattering experiments [4].

*In vivo*, repressor-mediated looping can be detected by monitoring transcription from a promoter that is controlled by two operators. Changing the spacing between the two DNA binding sites rotates the binding sites around the DNA helix relative to each other. Thus, some spacings are better for tetramer binding – and have better repression – than others [38, 53-56]. Using a second *in vivo* assay (comprising different cells strains and operators), we previously tested looping for the parent LLhP and LLhG+K chimeras [2]. Assays were carried out in *E. coli* strains that contained modified *lac* operons under control of the engineered *lacO*^sym^ [14] and natural *lacO*^2^ operators [2, 55]. Consistent with the known tetramerization propensities of GalR and PurR, LLhG+K exhibited changes consistent with looping whereas LLhP did not (Fig 3D) [2].

Thus, we considered whether both the offset and the slope differences between LLhP and LLhG+K variants (Fig 2) could be explained by LLhG+K looping in the original *in vivo* repression assay. These assays were carried out in an *E. coli* strain that contained a nearly wild-type *lac* operon (only *lacI* was interrupted). This operon comprises multiple DNA operators [57] – *lacO*^1^, *lacO^2^*, and *lacO^3^* (Fig 3; Table 1). Of these three natural operators, *lacO^1^* showed the highest affinity for wild-type LacI, *lacO^2^* exhibited 30-100 fold weaker binding (S1 Fig [19, 28, 58]), and *lacO^3^* binding was weaker still [15-18].

Following the example of wild-type LacI [20, 33, 38-41, 43, 44], LLhG+K looping two *lac* operators should lead to enhanced repression relative to non-looping LLhP, and thus the overall offset seen in Fig 2. The difference in LLhP and LLhG+K slopes (Fig 2) could be explained by changes in the local concentration of repressor that would coincide with altered *K_d_* for *lacO^1^* [59]: When tetramer stochastically dissociates from one of the two operator sites, binding to the other site would keep the repressor in the local vicinity, impeding competition by nonspecific genomic DNA. Thus, increasing affinity for *lacO^1^* would increase both the residence time at *lacO^1^* and the local concentration of repressor at auxiliary operators. This in turn would lead to the increased slope for LLhG+K relative to non-looping LLhP.

Next, we considered which of the two auxiliary operators (*lacO^2^* or *lacO^3^*) was most likely to contribute to *in vivo* repression. Based on the very weak binding of wild-type LacI to *lacO^3^* (S1 Fig, [16, 17, 44, 59]), we reasoned that *lacO^2^* was most likely to be involved in LLhG+K looping. Thus, equilibrium dissociation constants for this operator were determined using purified proteins and operator. The nanomolar binding affinities observed (Table 2; S2 Fig) are sufficiently strong to contribute to repression. However, among the nine LLhG+K variants assessed, *lacO^2^* binding showed at most ~4-fold change (Fig 4), which was a much narrower range than expected from *lacO^1^* measurements. Indeed, when correlated to binding affinities for *lacO^1^*, binding affinities for *lacO^2^* showed a slope that approached zero (Fig 4). Finally, for several variants, *lacO^2^* binding showed little effect from the addition of 10 mM fucose inducer (S2 Fig, closed squares).

**Figure 4.**
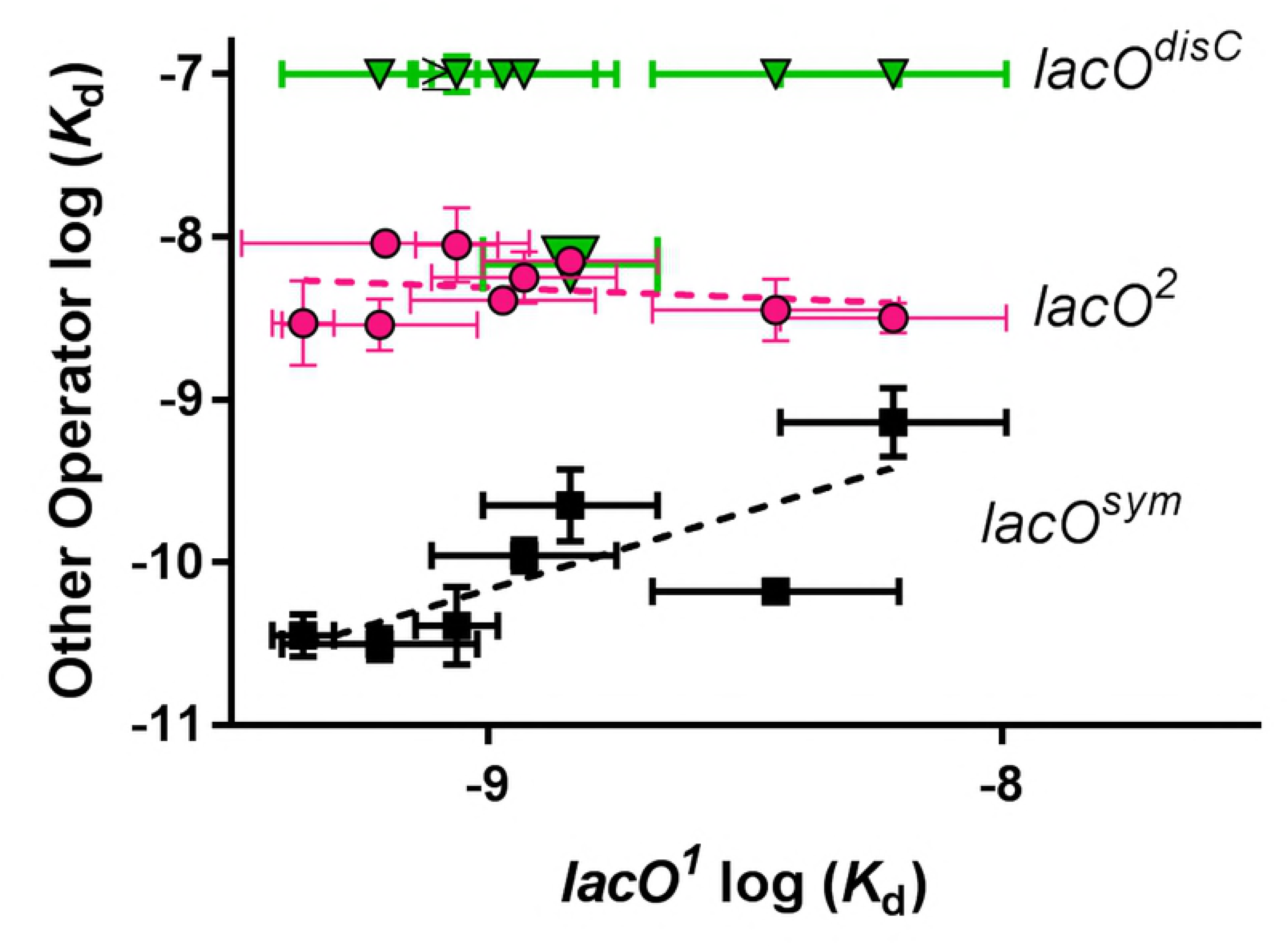
Comparison of *in vivo* repression and *in vitro* binding for LLhG+K variants binding to various operators. For LLhG+K variant proteins, the *K*_d_ values for binding to operators *lacO*^2^ (magenta circles),*lacO*^sym^ (black squares), and *lacO*disC (green triangles) are plotted against *K*_d_ values for binding to operator *lacO^1^*. The large green triangle highlights the V52P variant that had tighter *lacO^disC^* binding than the other variants. *K*_d_ values for LLhG+K binding operator *lacO^1^* are from [3]. *K*_d_ values for *lacO^sym^* and *lacO^2^* are summarized in Table 2. For *lacO^disC^*, most *K*_d_ values were out of range for the binding assay and a lower limit is shown. The lines are to aid visual inspection of the data. Error bars on both the X and Y parameters represent one standard deviation of the average values.

These findings were unexpected and led us to wonder how amino acid changes among the LLhG+K variants altered binding to other operators, such as the engineered operators *lacO^sym^* and *lacO*^disC^ (Table 1; S3 Fig and S4 Fig). Binding to these operators was previously characterized for both LacI and LLhP variants (Fig 5). Most proteins bound *lacO^sym^* more tightly: Variants of LacI bound *lacO^sym^* up to 10-fold more tightly than *lacO^1^* [28, 58], as did five LLhP variants [4]; however, two LLhP variants exhibited very poor binding (*K_d_* >10^−7^ M) [4]. For *lacO^disC^*, most LacI variants bound ~100-fold more weakly than *lacO^1^* [19, 58], whereas LLhP variants bound *lacO^disC^* 5-10-fold more weakly than *lacO^1^* [4].

**Figure 5.**
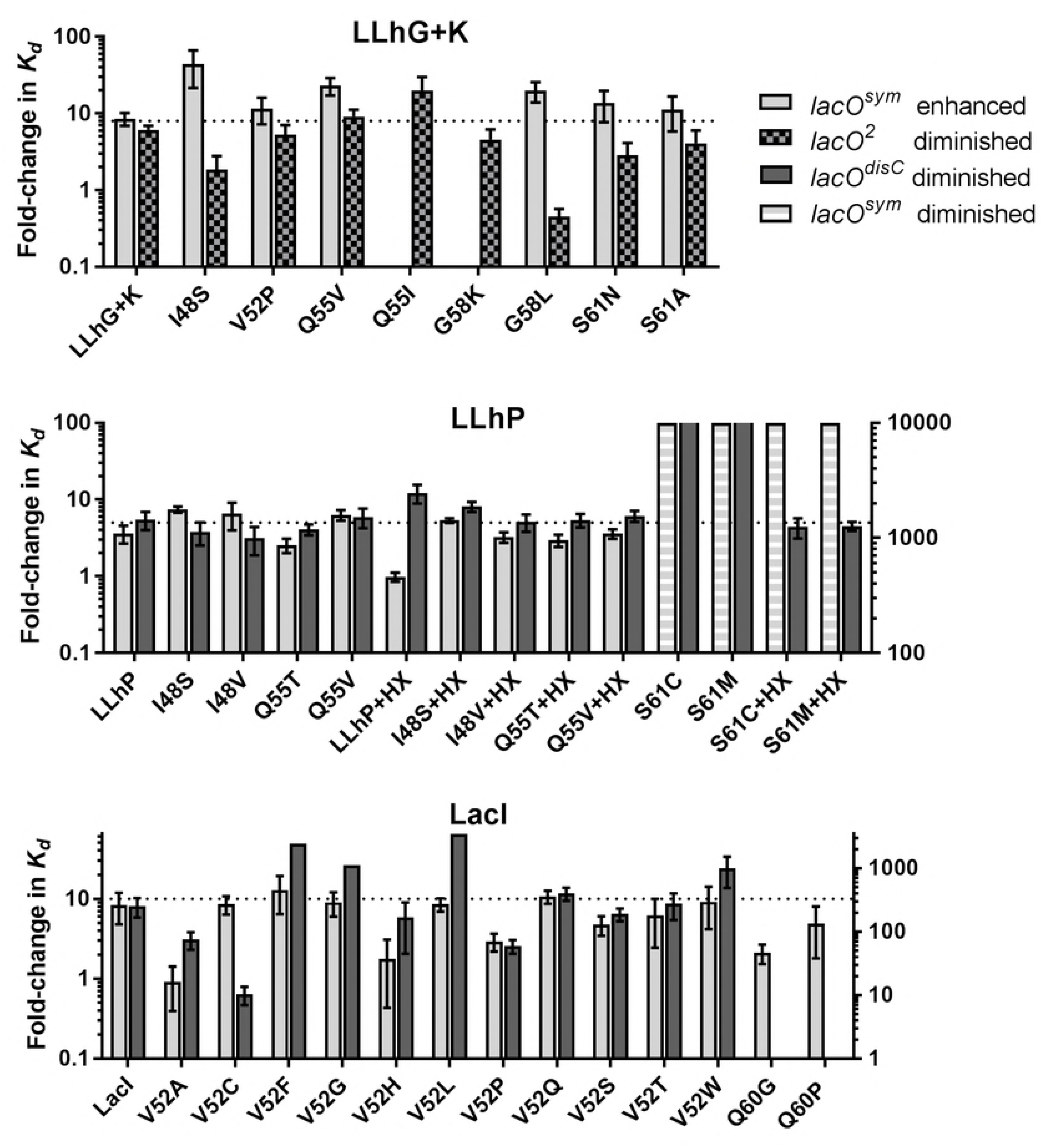
Altered fold-change in operator binding indicates altered DNA specificity. Fold-change for binding to the indicated operators was calculated relative to the *K_d_* for binding *lacO^1^*. To aid recognition of specificity changes, the ranges of the left and right y axes were chosen so that fold-change for *lacO^sym^* (left y axis) and *lacO^2^* (LLhG+K, left y axis) and *lacO^disC^* (LLhP and LacI, right y axis) were visually similar for the parent proteins. For LLhP variants at position 61, *lacO^sym^* binding was also diminished and is also plotted on the right y axis. Error bars were propagated from the standard deviations of average *K_d_* values reported in this manuscript and in previous publications [3, 4, 28, 58]. The dotted line is to aid visual comparison of the parent proteins with their amino acid variants. For each variant, (i) if fold-change for one operator deviates from the dotted line, or (ii) if fold-change of the two bars deviate from each other, then the DNA specificity of the variant has changed relative to the parent repressor protein.

Analogous to results for LacI and LLhP, LLhG+K variants binding to *lacO^sym^* was enhanced and responded to inducer (Fig 4 black squares; S3 Fig; Table 2). The increase in *lacO^sym^* binding over *lacO^1^* binding was not perfectly uniform (*i.e.* scatter observed for Fig 4 black squares). Nevertheless, mutational outcomes for *lacO^sym^* and *lacO^1^* were much better correlated (slope approaching 1; Fig 4, black dashed line) than those for *lacO^2^* and *lacO^1^* (slope near zero; Fig 4, magenta dashed line). For *lacO^disC^*, binding was above the limit of the filter binding assay for most LLhG+K variants (Fig 4; S4 Fig), which was a much larger fold-change than previously observed for LLhP variants. Nevertheless, LLhG+K V52P had measurable binding to *lacO*^disC^C (Fig 4; S4 Fig). Although it seems surprising that a proline in the middle of the helix (Fig 1B) allowed DNA binding, a similar outcome was observed for V52P in wild-type LacI [28].

## Discussion

*In vivo* activity is usually the sum of many protein activities. In our attempts to dissect the parameters relevant to *in vivo* repression of the Lac-based transcription repressors, we unexpectedly discovered that – while amino acid changes in LLhG+K did alter *lacO^1^* and *lacO^sym^* binding – they had very little impact on *lacO^2^* binding (Fig 4). This phenomenon was not simply a property of weaker binding for LLhG+K and *lacO^2^*: LLhP variant binding to *lacO^1^* and *lacO^disC^* spanned a similar magnitude yet showed the expected sensitivity to amino acid variation [4].

These results raise the question as to how these outcome is expected to generalize to other LLhG+K variants or to other LacI/GalR homologs. The amino acid changes in the current study were located throughout the LLhG+K linker structure (Fig 1B); thus, we expect that *lacO^2^* binding may generally lack sensitivity to changes in this region of this protein. However, whether *lacO^2^* mutational insensitivity is unique to LLhG+K or a general property of any LacI-based repressor remains to be seen. Such studies have not been carried out even for variants of full-length LacI, and the three homologs and their variants studied to date have enough differences (Fig 5 and discussed further below) to preclude extrapolating binding behaviors from one protein.

Another consideration raised by the current results is the comparison of *lacO^2^* binding to nonspecific binding. LLhG+K binding to *lacO^2^*, with its similar binding affinities of variants/lack of induction, is reminiscent of LacI binding to non-specific (genomic) DNA [30]. However, LLhG+K binding affinities were up to five orders of magnitude tighter than expected for non-specific binding, which is estimated to be 3 x 10^−4^ M for wild-type LacI [29]. Furthermore, non-induction is not a general property of *lacO^2^*, since wild-type LacI binding to *lacO^2^* was diminished in the presence of IPTG (S1 Fig). More experiments would be required to assess non-specific binding by LLhG+K. Likewise, although LacI binding to *lacO^3^* was much weaker than the detection limits of the assay used in the current study (S1 Fig) and thus not pursued, some LLhG+K variants might have unexpected interactions with *lacO^3^*.

These results raise several points that should be kept in mind when constructing synthetic transcription circuits. First, one should be aware whether or not alternate operators are present. If the LacI/LacZ combination is used as the reporter protein for circuit development, *lacO^2^* will naturally present at the start of the *lacZ* gene [16]. (Since remnants of the *lacZ* gene might also contain the *lacO^2^* operator sequence, discrepancies could arise even if another reporter gene is used.) Second, in fine-tuning circuits for desired output, one could mutate the operator sequence to alter baseline or induced expression levels. If, for example, a multi-input circuit was built using LacI-based chimeras (*e.g.* [1]) and the operator sequence was changed to reduce baseline expression, one should not assume that the repressor-operator interaction will be equally altered for all chimeras. Third, we expect this phenomenon could be observed for broad range of transcription factors that bind to alternative engineered or natural operator sequences.

More broadly, these results lead us to look at the criteria for quantitatively assessing ligand specificity changes. We previously used the rank order of ligand affinities to assess whether changes in the region altered ligand specificity [4, 60]. The current work shows that this definition was too narrow. In his seminal textbook, Creighton stated “Specific binding by a protein of one ligand, and not another, depends on their relative affinities, their concentrations, and whether they bind at the same site” [61]. By this definition, a specificity change would also be indicated by differences in the fold-change among ligands, even if the rank order stayed the same. Interestingly, fold-change among variant operators was similar for most LLhP variants studied, in contrast to the fold-changes differences observed among the LLhG+K variants and variants at LacI position 52 (Fig 5) [3, 4, 28]. Thus, this comparison provides another example for which the functional attributes of one protein cannot be extrapolated to other family members.

This extrapolation limitation is especially relevant when considering algorithms that predict ligand specificity from sequence alignments. Indeed, the linker positions mutated in this study were predicted to be specificity determinants (that is, locations that can be substituted to alter specificity) for the naturally occurring LacI/GalR homologs (discussed in [62]). We previously concluded from the LLhP studies that changes at these linker positions affected overall binding affinity more often than specificity. However, in LLhG+K, variants at linker positions show fold-change differences indicative of specificity changes (Fig 5). Perhaps our LLhP studies were too limited in scope to detect specificity changes. Alternatively, one unified set of “specificity determinants” may not be appropriate for defining ligand specificity across the whole family. This conclusion is consistent with previous analyses of individual LacI/GalR subfamilies, which predicted that the locations of positions important to each subfamily fall in different places on the common LacI/GalR structure [63].

The complexity of the observed specificity changes may be analogous to the non-additive outcomes that often arise when multiple amino acids are substituted in one protein (epistasis). In the LacI-based repressors, we noted considerable epistasis arose from combinatorial changes in the linker region [11, 12]. Ligand variation could be thought of as one more mechanism for changing the chemical environment that, in turn, alters the outcome of chemical changes that accompany amino acid substitution.

## Acknowledgements

We thank Ms. Edina Kosa for assistance with DNA binding assays and Dr. Sarah Bondos (Texas A&M Health Science Center) for discussions about DNA binding and comments on the manuscript. We thank Drs. Ernesto Fuentes (University of Iowa), Brian Baker (Notre Dame University), and Marina Ramirez-Alvaredo (Mayo College of Medicine) for helpful discussions about the definition of “ligand specificity”.

**Supporting Information**

**S1 Fig.** Representative curves for LacI binding to *lacO^2^* and *lacO^3^* operators.

**S2 Fig.** Representative curves for LLhG+K variants binding to operator *lacO*^2^.

**S3 Fig.** Representative curves for LLhG+K variants binding to operator *lacO*sym.

**S4 Fig.** Representative curves for LLhG+K variants binding to operator *lacO^disC^*.

**S5 Fig.** Representative curve for LLhG+K S61A binding to operator *lacO*^1^.

